# Modelling early thermal injury using an *ex vivo* human skin model of contact burns

**DOI:** 10.1101/2020.08.18.254458

**Authors:** Aiping Liu, Edgar Ocotl, Aos Karim, Josiah J. Wolf, Benjamin L. Cox, Kevin W. Eliceiri, Angela LF Gibson

## Abstract

**Background:** Early mechanisms underlying the progressive tissue death and the regenerative capability of burn wounds are understudied in human skin. A clinically relevant, reproducible model for human burn wound healing is needed to elucidate the early changes in the human burn wound environment. This study reports a reproducible contact burn model on human skin that explores the extent of tissue injury and healing over time, and defines the inter-individual variability in human skin to enable use in mechanistic studies on burn wound progression and healing.

**Methods:** Using a customized burn device, contact burns of various depths were created on human skin by two operators and were evaluated for histologic depth by three raters to determine reproducibility. Early burn wound progression and wound healing were also evaluated histologically after the thermally injured human skin was cultured *ex vivo* for up to 14 days.

**Results:** Burn depths were reproducibly generated on human skin in a temperature- or time-dependent manner. No significant difference in operator-created or rater-determined depth was observed within each patient sample. However, significant inter-individual variation was identified in burn depth in ten patient samples. Burn-injured *ex vivo* human skin placed into culture demonstrated differential progression of cell death and collagen denaturation for high and low temperature contact burns, while re-epithelialization was observed in superficial burn wounds over a period of 14 days.

**Conclusion:** This model represents an invaluable tool to evaluate the inter-individual variability in early burn wound progression and wound healing to complement current animal models and enhance the translation of preclinical research to improvements in patient care.

## 1. Introduction

The depth of a burn injury in skin determines the healing capacity of the wound and the need for surgical intervention [1]. With recent advancements in tissue engineering, the ability to support wound healing in deep partial thickness burn injury may mean that fewer patients will require the creation of a wound (donor site) to surgically treat a burn wound. However, the early stages of burn wound “conversion,” a process by which burns progress to a deeper injury, have been shown in multiple animal models but remain understudied in human skin [2, 3]. There is a dearth of knowledge about the early events that occur in the burn microenvironment. These events may represent unique targets for intervention to enhance wound healing in human skin.

Animal models have been used extensively to study burn wound healing. While rodents are the most widely used small-scale animal models in burn research due to ease of maintenance, fast breeding, and transgenic strain generation in laboratory settings [4], researchers have become increasingly aware of the potential limitations of extrapolating data from rodents to humans [5]. The most closely related animals for modelling human burn injury are pig, which have many similarities in skin anatomy. Whereas rodent wound healing occurs by contraction [5], wounds in pig and humans heal by re-epithelialization [6–8]. Despite the similarities to human skin, differences exist that may impact the translation of findings in pig to humans [7, 9]. Additionally, ethical issues, high cost, and cumbersome handling may limit the ability to use this model in a high throughput manner. *In vitro* models based on monolayer cell culture (mouse or human cells) are the simplest system to characterize cellular signaling events in response to thermal injury [10, 11]. To mimic the three-dimensional environment of cells, human keratinocytes have also been seeded onto a de-epidermized dermis and cultured up to 20 days to study the cell-matrix interaction and cell migration in burn research [12]. Although indispensable for dissecting the detailed molecular and cellular mechanisms in burn injury, these *in vitro* models lack the complex architecture of human skin.

Because an *ex vivo* model of human skin maintains physiological skin biology, skin barrier function, and metabolism during culture [13], it has been suggested as the “missing link” that bridges *in vitro* models of wound healing and clinical practice. For burn wounds, several *ex vivo* human skin models have been developed to comprehensively characterize the response of cells and extracellular matrix to the thermal injury and early stages of healing processes [14–16]. However, these models of human skin have not thoroughly described the time and temperature relationship of thermal injury, and the inter-individual variability that may exist. Here, we present a reproducible, contact burn model on *ex vivo* human skin. The model replicates the extent of burn injury and healing over time that is observed clinically. In addition, we highlight the inter-individual variability in thermally injured human skin. This high-throughput, inexpensive model has wide application for researchers investigating pathology, novel burn therapies, or diagnostics for burn injury.

## 2. Materials and Methods

### 2.1. Human Skin Tissue Preparation

Human skin was obtained from patients undergoing elective reconstructive surgeries at our institution. The de-identified samples were exempt from the regulation of University of Wisconsin-Madison Human Subjects Committee Institutional Review Boards. Data on patient age, sex, and type of surgery were collected with the tissues. The skin samples were processed within an hour after surgery. Excess subcutaneous fat was removed from the tissue to make the skin surface even. At the time of burn, the initial temperature of the tissue was recorded and was at the room temperature (unless otherwise specified).

### 2.2. Burn Device

The device used to burn human skin samples was customized and fabricated (Fab Lab, Morgridge Institute for Research, Madison, Wisconsin). The 8mm circular burn surface was produced from a stainless-steel cylinder that was partially hollowed and press-fit onto a lathe-modified tip of a mini soldering iron (TS100, GtFPV, FL). The stainless-steel cylinder was electronically heated, via the internal soldering iron, by setting the soldering iron to a desired user-preset temperature that ranges between 100°C to 150°C.The soldering iron itself was smoothly delivered to contact tissue samples by attachment to a small, vertically oriented ball-bearing carriage and guiderail to create uniform, 8mm circular burn wounds (Fig. 1). The actual operating temperature at the burn tip surface, which is slightly lower (<1 °C) than the preset temperature due to the heat loss through the cylinder, was monitored in real-time throughout burn wound creation by attaching a K-type thermocouple (SA1-K-SC, Omega, CT) and a digital multi-meter (TM-902C, Lutron Electronics, PA). The weight of the cylindrical stainless-steel block provided a standardized pressure (40g) that was applied to the human skin during burn creation. This burn device has been provided as an open-source tool; all relevant part files, as well as a more detailed description of the device itself, are available for download at: https://www.morgridge.org/designs.

**Fig. 1.**
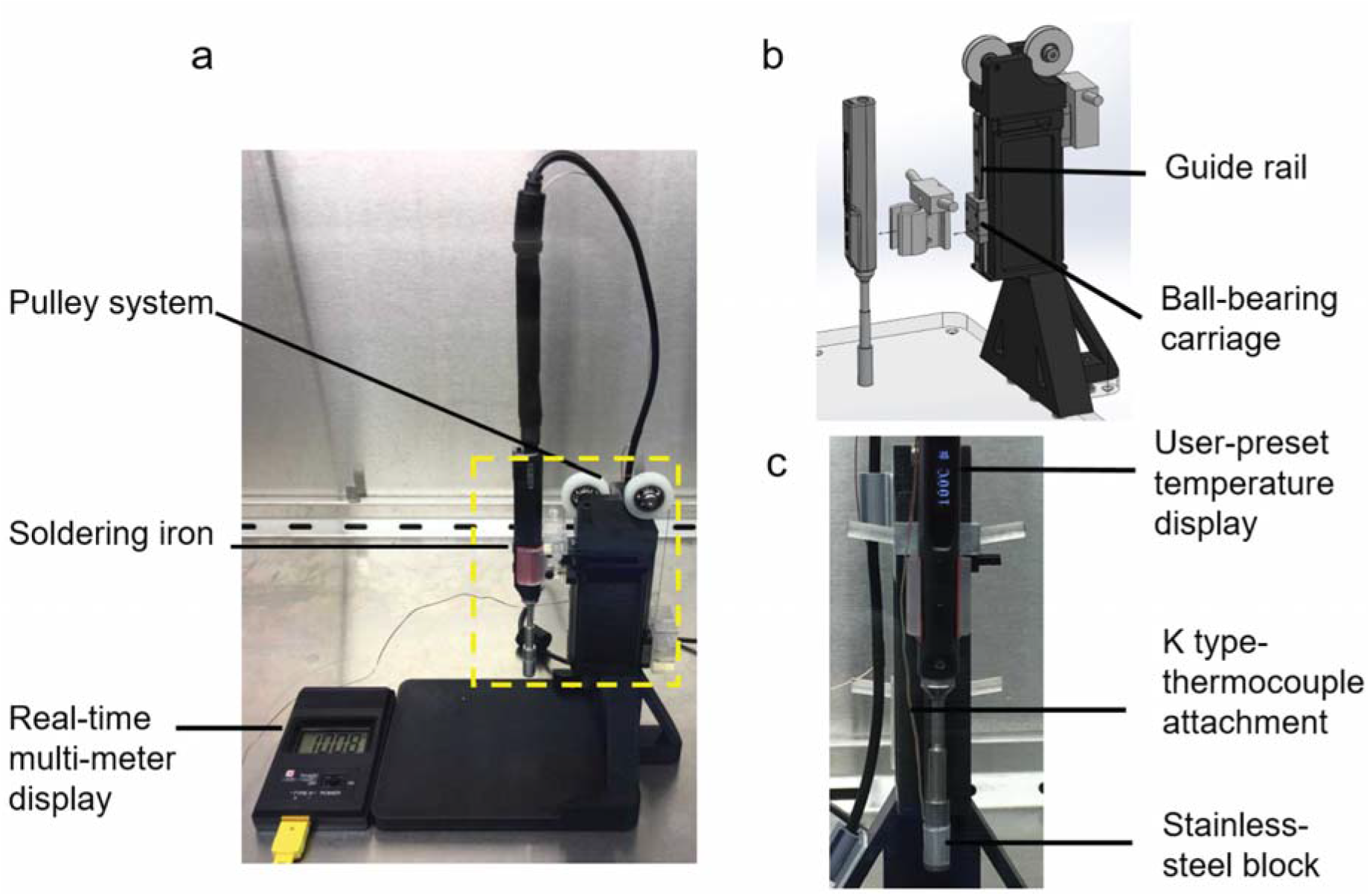
Customized contact burn device. (a) Side view of the full assembly of the burn device. (b) Exploded view of the pulley system, showing the soldering iron attached to the vertically-oriented ball bearing carriage and guiderail. (c) Front view of the soldering iron.

### 2.3. Experimental Protocols for Model Characterization

We conducted a series of experiments (n = 6) with different goals (see Table 1). For experiments 1-3, 5, and 6, each experiment used skin tissue from a single donor with three technical replicates for each burn condition (i.e., different temperatures or burn durations). Experiment 4 used skin tissue from 10 donors. For each donor tissue-a total of 6 biopsies were used to obtain 3 technical replicates per operator. All tissue was obtained from the abdomen. Burns were created on human skin tissues using the customized burn device and the parameters listed in Table 1. Briefly, a large piece of de-identified human skin tissue was obtained from plastic surgery procedures using an Institutional Review Board exempt protocol in accordance with laws and regulations of the University of Wisconsin-Madison School of Medicine and Public Health. The skin tissue was tacked down to foam on a slight stretch to create a uniform stabilized surface on the skin prior to thermal injury. The burns were created on the larger piece of skin tissue prior to taking a biopsy of the injured skin for further analyses. For the studies on the pathophysiology between high and low temperature burns (Experiment 2) and burn wound healing over time (Experiment 3), tissue biopsies were placed into tissue culture after initial burn. To examine the pathophysiology or healing of the burns wounds, separate tissue biopsies (n = 3 at each time point) were cultured and processed for histological examination at specific time points, which were also noted in Table 1. For the other studies, tissue biopsies were processed immediately after burn.

**Table 1.**
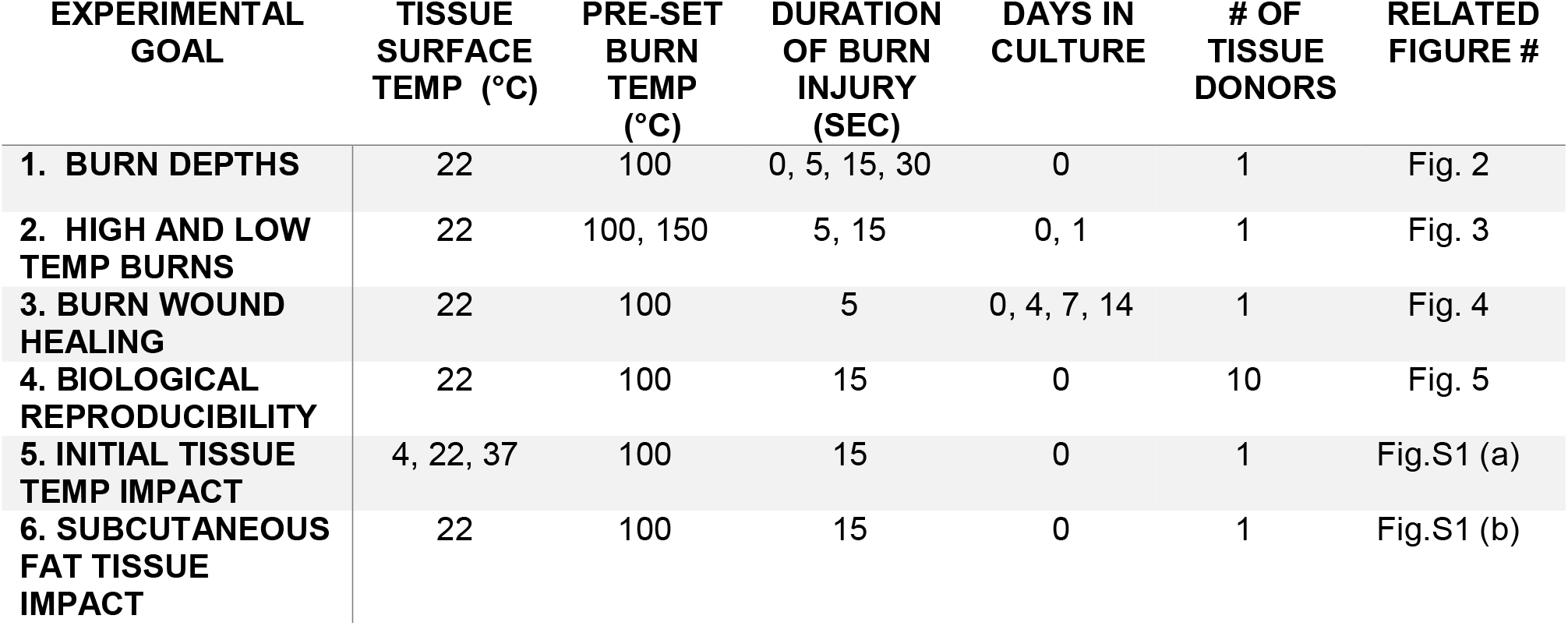
Burn creation parameters for model characterization.

| EXPERIMENTAL GOAL | TISSUE SURFACE TEMP (°C) | PRE-SET BURN TEMP (°C) | DURATION OF BURN INJURY (SEC) | DAYS IN CULTURE | # OF TISSUE DONORS | RELATED FIGURE # |
| --- | --- | --- | --- | --- | --- | --- |
| 1. BURN DEPTHS | 22 | 100 | 0, 5, 15, 30 | 0 | 1 | Fig. 2 |
| 2. HIGH AND LOW TEMP BURNS | 22 | 100, 150 | 5, 15 | 0, 1 | 1 | Fig. 3 |
| 3. BURN WOUND HEALING | 22 | 100 | 5 | 0, 4, 7, 14 | 1 | Fig. 4 |
| 4. BIOLOGICAL REPRODUCIBILITY | 22 | 100 | 15 | 0 | 10 | Fig. 5 |
| 5. INITIAL TISSUE TEMP IMPACT | 4, 22, 37 | 100 | 15 | 0 | 1 | Fig.S1 (a) |
| 6. SUBCUTANEOUS FAT TISSUE IMPACT | 22 | 100 | 15 | 0 | 1 | Fig.S1 (b) |

### 2.4. Tissue Culture

After thermal injury was induced, full thickness biopsies of human skin samples were taken by a 12-mm biopsy punch with the area of burned tissue (8 mm) in the center with surrounding normal unburned skin. Biopsies were placed on a customized, elevated metal mesh insert in p100 culture dish, which allowed the biopsies to be cultured at the air-liquid interface in culture media. Culture media contained DMEM (Gibco Laboratories, Gaithersburg, MD) supplemented with 10% FBS (Hyclone Laboratories Inc, Logan, UT), 0.625 μg/ml Amphotericin B (Gemini Bio, Sacramento, CA), and 100 μg/ml of Pen/Strep (Gibco Laboratories, Gaithersburg, MD) and was changed every other day to prevent exhaustion of nutrients [17]. For the wound healing studies, culture media were further supplemented with 10 μg/mL of insulin (Sigma-Aldrich, St. Louis, MO), 10 ng/mL of hydrocortisone (Sigma-Aldrich, St. Louis, MO), and 2 mM of L-glutamine (Sigma-Aldrich, St. Louis, MO) [18]. These samples were kept in an incubator at 37 °C and 5% CO_2_ for up to 14 days.

### 2.5. Histological Tissue Processing

Tissue biopsies were bisected, and one half was cryopreserved with Tissue-Tek® optimum cutting temperature (OCT) compound (Sakura Finetek USA Inc, Torrance), while the other half was fixed in 10% neural buffered formalin for paraffin embedding. The OCT frozen samples were cryo-sectioned into 8 μm thickness slides and stained for lactate dehydrogenase (LDH) [19] and Elastin Van Gieson (EVG) [20]. Routine hematoxylin and eosin (H&E) staining was performed on 5 μm thickness paraffin sections.

### 2.6. Burn Wound Assessment

Tissue sections were examined using a Nikon Ti-S inverted microscope. Images were captured using Nikon DS Ri2 Cooled color camera and Nikon imaging software, NIS Elements (Nikon Instruments Inc, Melville, NY). The primary outcome measured was the depth of thermal injury based on two dermal components: cellular viability and collagen injury. The depth of collagen injury was determined by measuring the vertical distance between the cutaneous basement membrane and the deepest level of magenta discoloration in EVG stained sections. The depth of cellular injury was determined by measuring the vertical distance between the cutaneous basement membrane and the initial depth where there was blue staining in epithelial, fibroblast, immune and endothelial cells in LDH stained sections. The relative burn depth was calculated for each individual burn by dividing the absolute depth of injury by the total thickness of the dermis. Burn depth was scored based on cell viability (LDH slides) using a score system (Fig. 2 e) modified from previous studies [19]. To assess the technical variation resulting from different operators using the burn device and rater bias on histological assessment, we compared the burn injuries of ten patients from two operators and the scores from three raters. The re-epithelialization of burn wounds was evaluated qualitatively on H&E stained tissue sections.

**Fig. 2.**
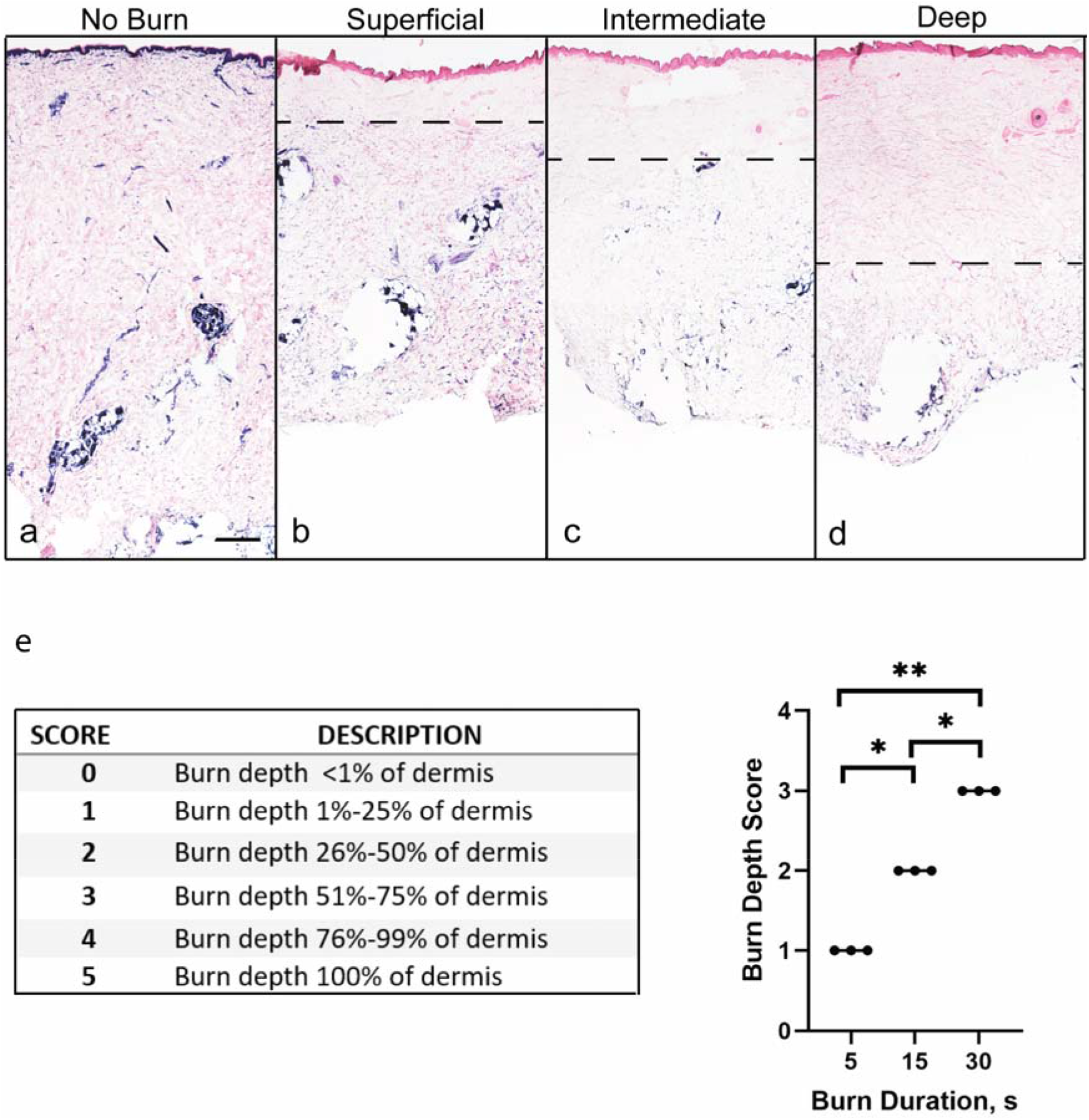
Spectrum of burn depths created through varying contact times. LDH staining (blue = viable cells) of (a) normal human skin without burn, and skin burned at 100 °C for (b) 5 seconds, (c) 15 s, and (d) 30 s. The burn depth into the dermis is represented by a dashed line. The relative burn depth in each section was quantified as the percentage of dermis lacking cell viability normalized to the total dermal thickness; scores of 0-5 were assigned as illustrated in (e). Note that tissue compression occurs in the burned samples (b-d) compared with non-burned sample (a). LDH stained sections are representative samples of 3 technical replicates per burn duration. Scale bar: 500 μm. One-way ANOVA and Tukey’s multiple comparison test was performed on the relative burn depth. *, P<0.05, **, P<0.005.

### 2.7. Statistical Analysis

The data are presented as mean ± standard deviation. Statistical significance among the relative burn depths at different burn durations was analyzed with one-way analysis of variance (ANOVA) and Tukey’s multiple comparisons test. Biological variation in burn creation between patients was tested using two-way repeated measures ANOVA. A Spearman’s correlation was used to calculate the major factors that contribute to biological variation in histological scoring of dermal damage. Statistical analysis was performed using GraphPad Prims 8 software (GraphPad Software Inc, La Jolla, CA). p<0.05 was considered statistically significant.

## 3. Results

### 3.1. Burn injury depth is time-dependent

A customized burn device equipped with a pulley system was developed to enhance reproducibility of burn injury in the laboratory (Fig. 1). With the temperature of the burn device set @ 100 °C, contact times of 5, 15, or 30 seconds were used to generate burn injury depths on human skin, simulating clinically relevant superficial, partial, or deep partial thickness injury (Fig. 2 a-d). Increasing loss of cellular viability, evaluated immediately after burn injury, was demonstrated in a statistically significant linear relationship with increasing duration of contact (Fig. 2 e).

### 3.2. Burn wound progression is temperature-dependent

In addition to contact time, it is well understood that temperature also affects the depth of injury. However, the impact of temperature on cellular viability and the associated collagen denaturation is not well defined. Human skin was burned at two different time/temperature combinations to understand if there is a differential effect of temperature on cells and collagen (Fig. 3). Both low temperature–long contact time (100°C for 15 seconds) and high temperature–short contact time (150°C for 5 seconds) created superficial partial thickness burns on immediate assessment, affecting cell viability (Fig. 3 e, i) and collagen denaturation to similar depths (Fig. 3 f, j). After 1 day in tissue culture, the loss of cell viability progressed to deep partial thickness in both low and high temperature burns (Fig. 3 g, k). No change in cell viability or collagen was detected in the no burn samples (Fig. 3 a-d). Interestingly, the increase in the depth of collagen denaturation over time was only observed in high temperature burns (Fig. 3 h, l).

**Fig. 3.**
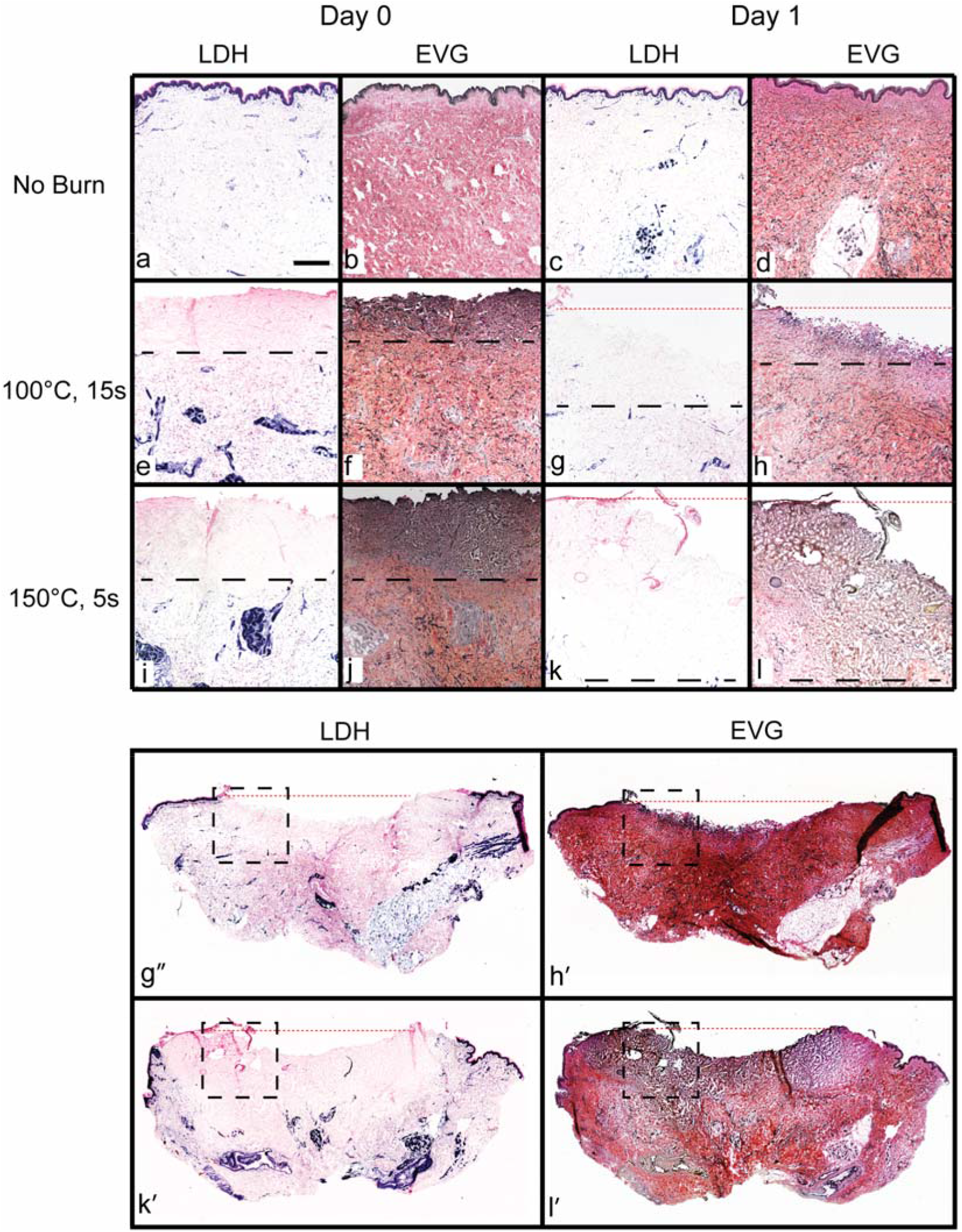
Effect of low and high temperature burns on cells and collagen structure diverges with time. Progressive cell death (black dash line in g, k) was observed for burns created at 100 °C for 15 s and 150 °C for 5 s, whereas the deepest level of progressive collagen damage (black dash line in h, l) was observed only in burns created at 150°C for 5 s. The red dotted line in g, h, k, or l indicates the superficial surface of the skin tissue, which was disrupted during tissue processing as a result of compromised integrity of burn tissues. g’, h’, k’ and l’ demonstrate the LDH and EVG stains of the entire tissue biopsies on Day 1 of culture with the dash box showing the location of g, h, k or l. Representative tissue sections of triplicate samples. Blue = viable cell in LDH stain; dark purple = denatured collagen in EVG stain. Scale bar: 500 μm.

### 3.3. Re-epithelialization occurs in *ex vivo* burn injured skin

In addition to illustrating the time/temperature association with burn injury depth and progression in human skin, this model can serve as a preclinical testing platform for burn wound healing. Burn wounds were created at 100°C for 15 seconds using our burn device. Tissue biopsies of the burn wounds were either processed for H&E staining immediately or after culturing for 4, 7 or 14 days. In Figure 4, we demonstrate re-epithelialization of a superficial partial thickness burn created *ex vivo* on human skin after 14 days in culture. These results are similar to re-epithelialization that occurs *in vivo* (Fig. 4 f). After 4 days in culture, reepithelization occurred from the wound edges (Fig. 4 c’), as well as from the hair follicles in the center of the wound (Fig. 4 c’’). The newly formed epidermal tongue proliferated and migrated underneath the damaged tissue and contained several organized layers of differentiating keratinocytes with a migrating tongue of epidermis. After 14 days of culture, neo-epidermis nearly covered the entire burn wound.

**Fig. 4.**
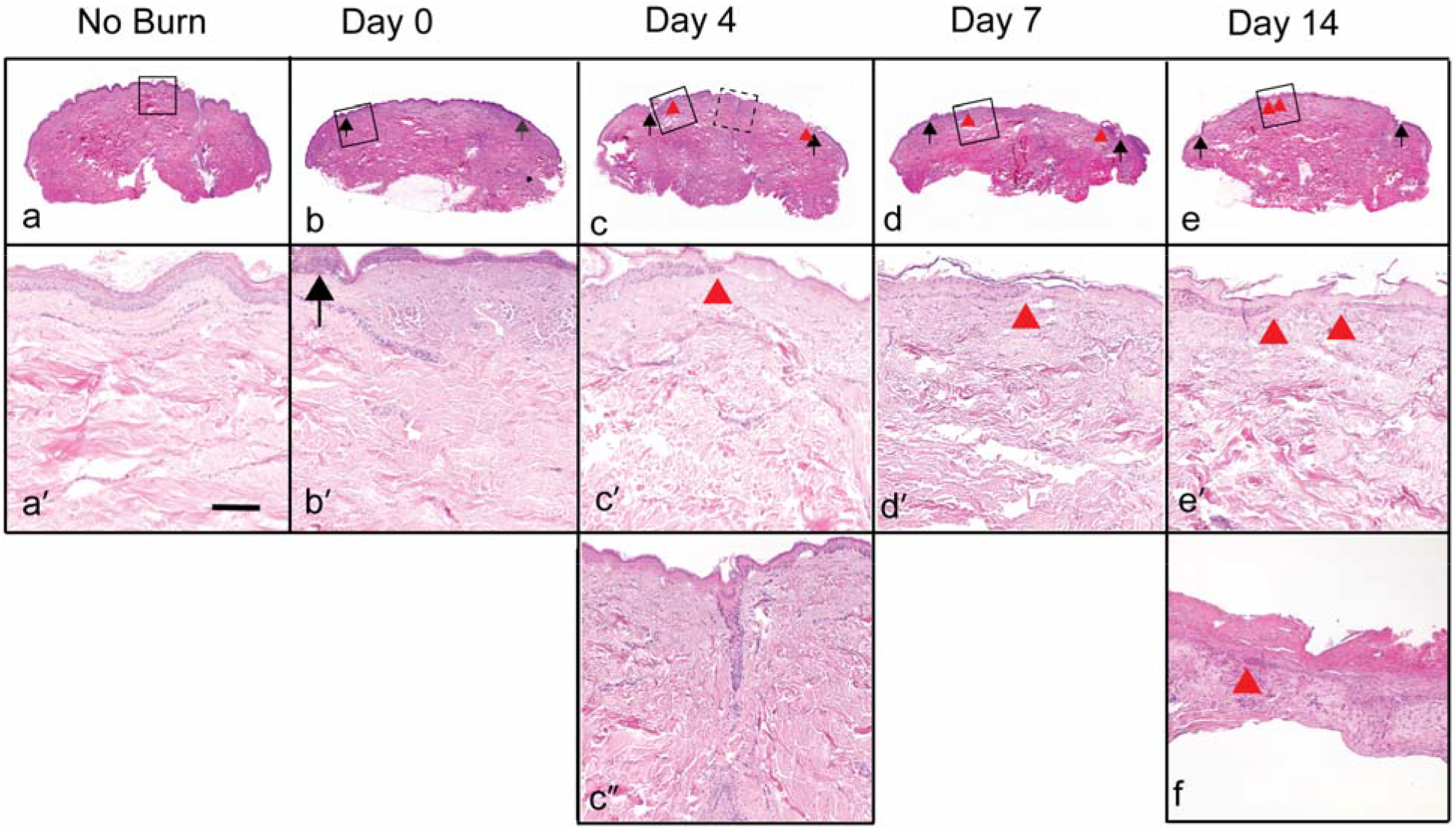
Re-epithelialization occurs from wound edges and dermal appendages after culture. H&E stained sections of normal human skin without burn (a, a’) and with burn immediately after injury (b, b’) or after (c, c’) 4, (d, d’) 7 and (e, e’) 14 days of culture. Black arrows indicate burn injured tissue margins and red arrow heads indicate the epithelial tongues of neo-epidermis over the burn. a’-e’ represent higher magnification of black inset boxes in a-e. (c”) Re-epithelialization emanating from a hair follicle in the burn wound in Day 4 (magnified image from dashed box in c). (f) H&E stained section of a tangential excision of burn wound tissue from a patient 14 days post burn injury, showing the epithelial tongue underneath the burned tissue slough. Scale bar: 200 μm. Representative images from triplicate samples.

### 3.4. Inter-individual variation exists in human skin burn injury

One advantage of animal models in wound healing research is that we can control the age, genetic makeup, and health of the animal. While this may be beneficial for reproducibility, the translatability of the preclinical research to humans, and the potential impact on society, are limited. We sought to investigate the degree of variation in burn depth after contact injury among 10 different patient skin specimens. The demographic data for each patient is listed in Table 2. Patients ranged in age from 27 years to 63 years, and the majority (80%) were female. Skin tissues from all 10 patients were subjected to the same burn condition (100°C for 15 seconds). We hypothesized that inter-individual variability exists in the time/temperature dependent effects of thermal contact and the resultant injury. No significant differences were observed in burn creation between two operators of the burn device (Fig. 5 a), however, a large variation (scores ranging from 1 to 5) in burn depths was observed between patient samples that received the same thermal insult, indicating that inter-individual variation exists. The rater subjectivity-dependent variance in scoring the LDH images for burn depth revealed no significant difference between raters (Fig. 5 b). Although a majority (> 60%) of the burns were scored as category 1 (1-25% burn depth) or category 2 (25-50% burn depth) with the same thermal insult, the burn depth scores vary significantly among patients, regardless of raters. To understand how patient demographics contribute to the variance of burn depths under the same thermal insult, we conducted Pearson R correlation among burn score, age, and skin thickness of the 10 patients. We found that age (R = 0.9) and skin thickness (R = 0.8) are significant contributors to the burn depth, with age positively and skin thickness negatively correlated to the burn depth, as would be expected.

**Fig. 5.**
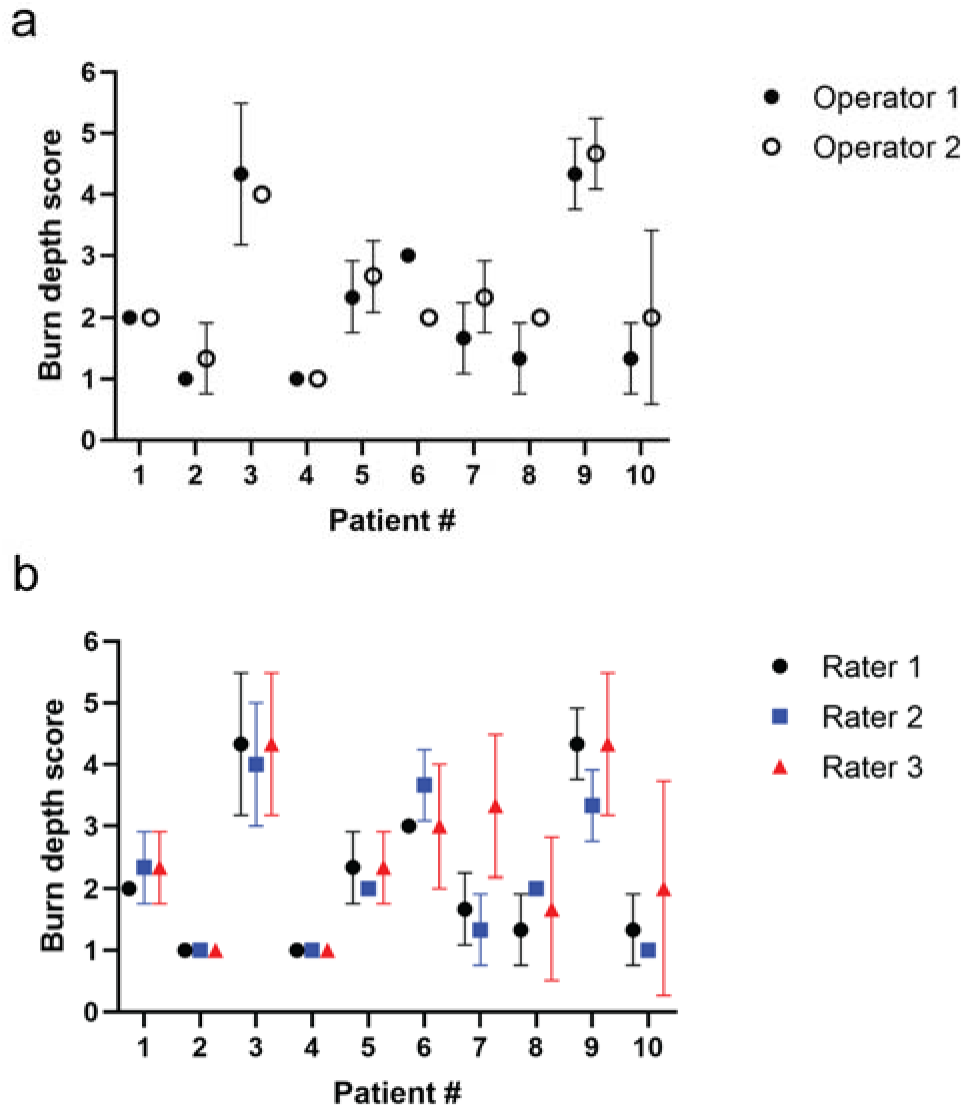
Biologic inter-individual variability in tissue injury depth after contact burn. (a) Operator-dependent variations in burn depth between two operators of burn device, (b) rater-dependent variations in burn depths among three raters. Data represents mean ± std with n = 3 samples per patient sample/operator/rater.

**Table 2.** Patient demographics and source of skin.

| **PATIENT**<br>**#** | **SEX** | **AGE** | **SURGERY** | **LOCATION** | **SKIN**<br>**THICKNESS**<br>**(MM) <sup>a</sup>** |
| --- | --- | --- | --- | --- | --- |
| 1 | Female | 40 | Abdominoplasty | Abdomen | 3.9±0.9 |
| 2 | Female | 35 | Bilateral DIEP | Abdomen | 5.5±1.0 |
| 3 | Female | 53 | Right DIEP | Abdomen | 3.9±0.5 |
| 4 | Male | 35 | Panniculectomy | Abdomen | 5.6±0.8 |
| 5 | Female | 47 | Right DIEP | Abdomen | 3.1±0.6 |
| 6 | Female | 52 | Abdominoplasty | Abdomen | 3.1±0.3 |
| 7 | Female | 53 | Abdominoplasty | Abdomen | 3.0±0.6 |
| 8 | Female | 33 | Abdominoplasty | Abdomen | 3.7±0.5 |
| 9 | Female | 63 | Left DIEP | Abdomen | 2.4±0.2 |
| 10 | Male | 27 | Abdominoplasty | Abdomen | 5.0±0.5 |

<sup>a</sup> Skin thickness data represents mean ± std with n = 6 samples per patient.

## 4. Discussion

The mechanisms of burn progression early after injury, and the minimum regenerative capacity that is necessary for healing without surgery in deep partial thickness burns, are fundamental questions that remain unknown [2, 3, 21]. The lack of a clinically relevant, reproducible human skin burn model is a major barrier to addressing these mechanistic and clinically important questions.

For burn models where a pre-defined and reproducible depth of injury is desired, a detailed investigation of all technical parameters in publications (temperature, pressure and exposure duration, or the actual amount of heat transferred) is critical to create reproducible or pre-defined burn depth [22–26]. In this study, we defined temperature, duration, and pressure exerted during burn creation on human skin (Table 1). We show consistent burn depths at a given temperature and duration of contact on human skin from a single donor (Fig. 2). By increasing the contact time, the injury depth increases in a linear relationship with the duration of contact, similar to porcine burn models [25, 27]. Other important variables for model development were identified during our studies. For example, initial tissue temperature and thickness of subcutaneous fat play a role in the depth of injury. Higher initial tissue temperature prior to creating the burn resulted in a deeper thermal injury (Supplemental Fig. 1 a). We chose room temperature as the initial tissue temperature because it is challenging to consistently maintain 37°C throughout the tissue during burn creation *ex vivo*. We also found that subcutaneous fat insulated the skin tissue against deeper injury with the same contact time/temperature combination (Supplemental Fig. 1 b), indicating a protective role of the fat tissue in cutaneous thermal injury. We left the subcutaneous fat attached to the tissue while performing the contact burns, with only minimal trimming to equalize the thickness of fat across the entire tissue to mimic the *in vivo* condition more closely.

Human skin tissue thickness and thermal properties vary with anatomic location, gender, and age [28–31]. These variations are predicted to have a pronounced influence on the extent of burn damage [32, 33]. To our knowledge, this is the first study to provide detailed histologic evidence of the biological inter-individual variations in human burn injury (Figure 5). These findings indicate that prior published burn depth estimates should be interpreted in the context of parameters such as anatomic location as it relates to skin thickness, initial tissue temperature, and method of injury (scald, contact, flame, etc) [34–36]. We feel these data illustrate the distribution of burn thresholds based on the diversity of human skin properties, and recommend increasing the sample size in studies to capture the expected variation. In addition, this work further confirms the misinterpretation of original work by Henriques and Moritz [36], which is propagated throughout the literature as discussed in an editorial by Abraham [37].

The variable temperature range setting of the burn device allows us to test whether differential damage occurs in cells and collagen depending on whether the skin is exposed to high or low temperature burns. Previous studies of low temperature contact burn (ranging between 60-90 °C) on pig skin have showed less depth of collagen injury compared with cellular injury [20, 22]. In this study, we found that both high (150 °C for 5 seconds) and low temperature (100°C for 15 seconds) burns have comparable initial damage in both cells and collagen (Fig. 3). However, after one day in culture the collagen damage in burns created at 100 °C appears more superficial than the cellular death, suggesting that the mechanism of burn wound progression in burns at different temperatures may differ, despite comparable initial injury patterns. In addition, we chose to use the EVG stain over Trichrome stain to identify collagen denaturation in this study because the EVG stain is able to detect more subtle changes in denatured collagen, thus it is more suitable for low temperature burns than Trichome stain (Supplemental Fig. 2).

There are several limitations of this study. First, the source of skin is limited to abdominal skin; therefore, we could not assess specific anatomic location-dependent effects of contact burn to depth of injury. Second, to minimize the variance in burn creation due to the difference in temperature on the surface and within the tissue as well as potential dissipation of heat over time, initial tissue temperature was set at the room temperature; consequently, the burn damage reported in this *ex vivo* model is likely less severe than that of *in vivo* burns from the same thermal insult. Third, although burn progression and re-epithelialization mimic the *in vivo* setting [27, 38, 39], the rates of progression and re-epithelialization depend on the composition of culture medium, which is physiologically different from that of an *in vivo* microenvironment. In addition, the tissue culture period was limited to 2 weeks, therefore observations of the time to fully re-epithelialize for all burns and final scar outcome were not possible. Most importantly, this model focuses on the contribution of local burn microenvironment without considering the contribution of systemic factors. Therefore, conclusions or recommendations drawn from this model should be cautiously interpreted with these limitations in mind and shall be supported by evidence from clinical observations in burn patients.

In conclusion, we have established a high-throughput, clinically relevant *ex vivo* human skin burn model that can be used for preclinical studies to further support findings in other animal models. The model is highly reproducible within a single donor tissue when parameters such as device pressure and temperature, initial skin temperature, and thickness of adipose tissue are controlled. Importantly, we found that inter-individual differences exist among tissue donors and should be taken into consideration when using human skin tissue for burn modelling. We believe this model will enhance the mechanistic evaluation of local mediators in the burn microenvironment that may be involved in burn progression, and early wound healing events that are otherwise challenging to study in human subjects *in vivo*.

## Abbreviations

EVG: Elastin van Gieson
DMEM: Dulbecco’s modified eagle medium
FBS: fetal bovine serum
H&E: hematoxylin and eosin
LDH: lactate dehydrogenase
OCT: Tissue-Tek^®^ optimum cutting temperature compound

## Compliance with ethical statements

### Ethical approval

This article does not contain any studies with human participants or animals performed by any of the authors. The human skin tissues used in this study were de-identified, therefore, exempted from approval of the Institutional Review Boards, University of Wisconsin-Madison.

## Acknowledgements

We acknowledge the assistance from the Fab Lab in the Morgridge Institute for Research, for computer-aided design and fabrication of our customized burn device, under funding from the BerbeeWalsh Prototype Pathway. We also acknowledge the work of statistical consultants Glen E. Leverson, PhD, and Lily N. Stalter, MS.

---

**Supplemental Fig. 1.**
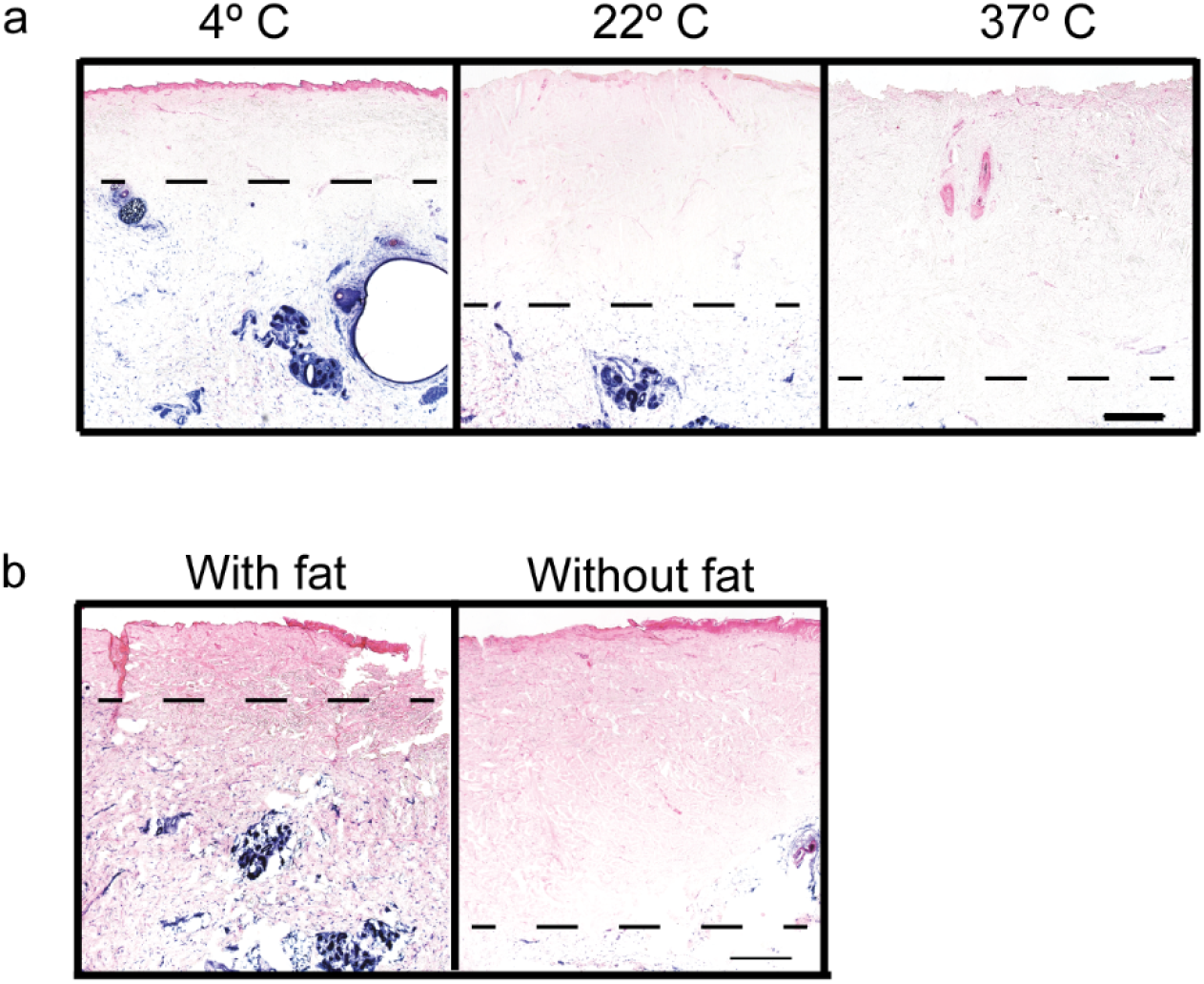
Impact of initial tissue temperature and fat tissue on burn creation on *ex vivo* human skin. Under the same contact burn settings, (a) the depth of burn injury increases with initial skin tissue temperature, and (b) burn depth increases in human skin when all subcutaneous fat tissue is removed. Scale bars: 500 μm.

**Supplemental Fig. 2.**
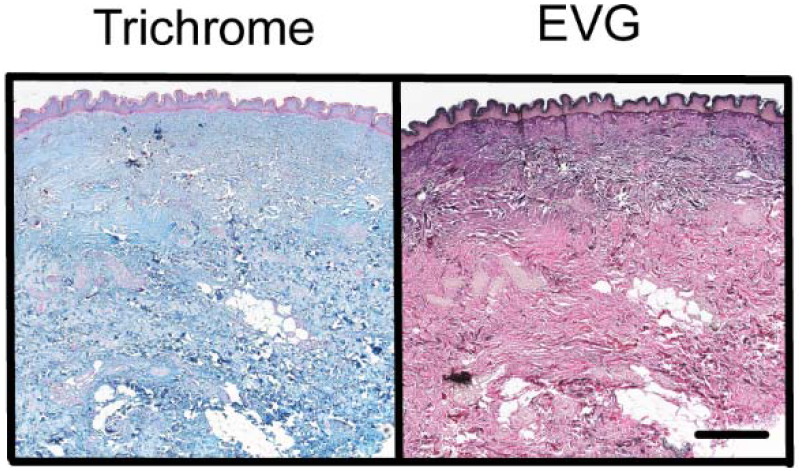
Comparison between trichrome and Elastin Van Gieson (EVG) stains for collagen denaturation in burn wounds. Collagen denaturation is demonstrated at the region of the purple color in the EVG stain but is not detected in the sequentially trichrome-stained section, suggesting that EVG is more sensitive for collagen denaturation in thermal burns (here, flame burn). Note, in Trichrome stain, denatured collagen is stained red, distinctive from intact collagen (blue), whereas in EVG stain, denature collagen is stained purple or dark blue in contrast to intact collagen, which appears red. Scale bar: 500 μm.

